# Neuroskeletal effects of chronic bioelectric nerve stimulation in health and diabetes

**DOI:** 10.1101/2020.11.24.396143

**Authors:** Alec T Beeve, Ivana Shen, Xiao Zhang, Kristann Magee, Ying Yan, Matthew MacEwan, Erica L Scheller

## Abstract

**Background/Aims:** Bioelectric nerve stimulation (eStim) is a novel clinical paradigm that can promote nerve regeneration after trauma, including within the context of diabetes. However, its ability to prevent the onset of diabetic peripheral neuropathy (DPN) has not yet been evaluated. Beyond the nerve itself, DPN has emerged as a potential contributor to sarcopenia and bone disease; thus, we hypothesized that eStim could serve as a strategy to simultaneously promote neural and musculoskeletal health in diabetes.

**Methods:** To address this question, an eStim paradigm pre-optimized to promote nerve regeneration was applied to the sciatic nerve, which directly innervates the tibia and lower limb, for 8-weeks in control and streptozotocin-induced type 1 diabetic (T1D) rats. Metabolic, gait, nerve and bone assessments were used to evaluate the progression of diabetes and the effect of sciatic nerve eStim on neuropathy and musculoskeletal disease, while also considering the effects of cuff placement and chronic eStim in otherwise healthy animals.

**Results:** Rats with T1D exhibited increased mechanical allodynia in the hindpaw, reduced muscle mass, decreased cortical and cancellous bone volume fraction (BVF), reduced cortical bone tissue mineral density (TMD), and decreased bone marrow adiposity. T1D also had an independent effect on gait. Placement of the cuff electrode alone was sufficient to alter gait patterns and to promote unilateral reductions in tibia length, cortical BVF, and bone marrow adiposity. Alterations in gait patterns and left-right balance to tibia length were restored with eStim, but it did not prevent T1D-induced changes in muscle, bone, marrow adiposity or mechanical sensitivity. Beyond this, chronic eStim resulted in an independent, bilateral reduction in cortical TMD.

**Conclusion:** Overall, these results provide new insight into the pathogenesis of diabetic neuroskeletal disease and its regulation by eStim. Though eStim did not prevent neural or musculoskeletal complications in T1D, our results demonstrate that clinical applications of peripheral neuromodulation ought to consider the impact of device placement and eStim on long-term skeletal health in both healthy individuals and those with metabolic disease. This includes monitoring for compounded bone loss to prevent unintended consequences including decreased bone mineral density and increased fracture risk.

## 1 Introduction

Bioelectric medicine has recently been gathering more interest as neuromodulation technologies advance and connections between the peripheral nervous system and target organs are better established. Technologies including fully implantable, self-powered systems (Yao et al., 2018) and bioresorbable electrodes with tunable degradation times (Koo et al., 2018) have expanded the possible applications of neuromodulation for long- and short-term use. In addition to applications in the central nervous system (Fishman et al., 2019; Lozano et al., 2019), bioelectric nerve stimulation (eStim) is now utilized in a variety of FDA-approved medical devices to treat peripheral disorders including chronic pain, obesity, and urinary incontinence (Reardon, 2014; McCrery et al., 2020). In the past two decades, eStim paradigms have also been established to enhance the regenerative capability of nerves in rodents and humans post-injury. In rats and mice, both motor and sensory nerves demonstrate upregulated regeneration-associated genes with application of eStim (Geremia et al., 2007; Brushart et al., 2005; Gordon and English, 2016). In humans, one application of post-surgical eStim similarly improves functional nerve regeneration after repair of digital nerve transection and median nerve crush injury (*i.e*. carpal tunnel syndrome) (Wong et al., 2015; Gordon et al., 2010)

Studies in mice and rats suggest that the neuroregenerative effects of eStim persist even in the metabolically challenged state of streptozotocin (STZ)-induced type 1 diabetes (T1D) (Singh et al., 2015; Lin et al., 2015). Therefore, we hypothesized that this paradigm may be sufficient to prevent the onset of diabetic peripheral neuropathy (DPN). Considering correlations between neural and musculoskeletal health in diabetes (Beeve et al., 2019; Forbes and Cooper, 2013; Jaiswal et al., 2017; Melendez-Ramirez et al., 2010), we also hypothesized that eStim would provide a simultaneous benefit for innervated downstream organs, including muscle and bone, in both control and diabetic rats. To test this hypothesis, we utilized a fully implantable, wireless system for sciatic nerve stimulation that has previously been employed to promote nerve regeneration (MacEwan et al., 2018). This technology enabled us to deliver weekly 1-hour eStim treatments for 8-weeks. Coupled with metabolic, neural and musculoskeletal analyses, our experimental design allowed us to determine the *in vivo* effect of chronic eStim on nerve, muscle and bone in the context of health and diabetes. This work was completed as part of the National Institutes of Health SPARC consortium (*<u>S</u>*timulating *<u>P</u>*eripheral *<u>A</u>*ctivity to *<u>R</u>*elieve *<u>C</u>*onditions) in the United States.

## 2 Materials and methods

### 2.1 eStim device fabrication

Sciatic nerve cuff stimulators were fabricated as previously described (MacEwan et al., 2018). Braided Pt/Ir leads (10IR9/49T, Medwire, Sigmund Cohn Corp.) were threaded through 8-mm of silicon tubing (inner diameter 1.5-mm) using a custom plastic rig. Exposed wires on the external surface of the silicon cuff were insulated with medical-grade silicone elastomer (A-564, Factor II). Cuff leads were then soldered to a thin-film wireless receiver coil and embedded in the same medical silicone (Fig.1A).

**Figure 1.**
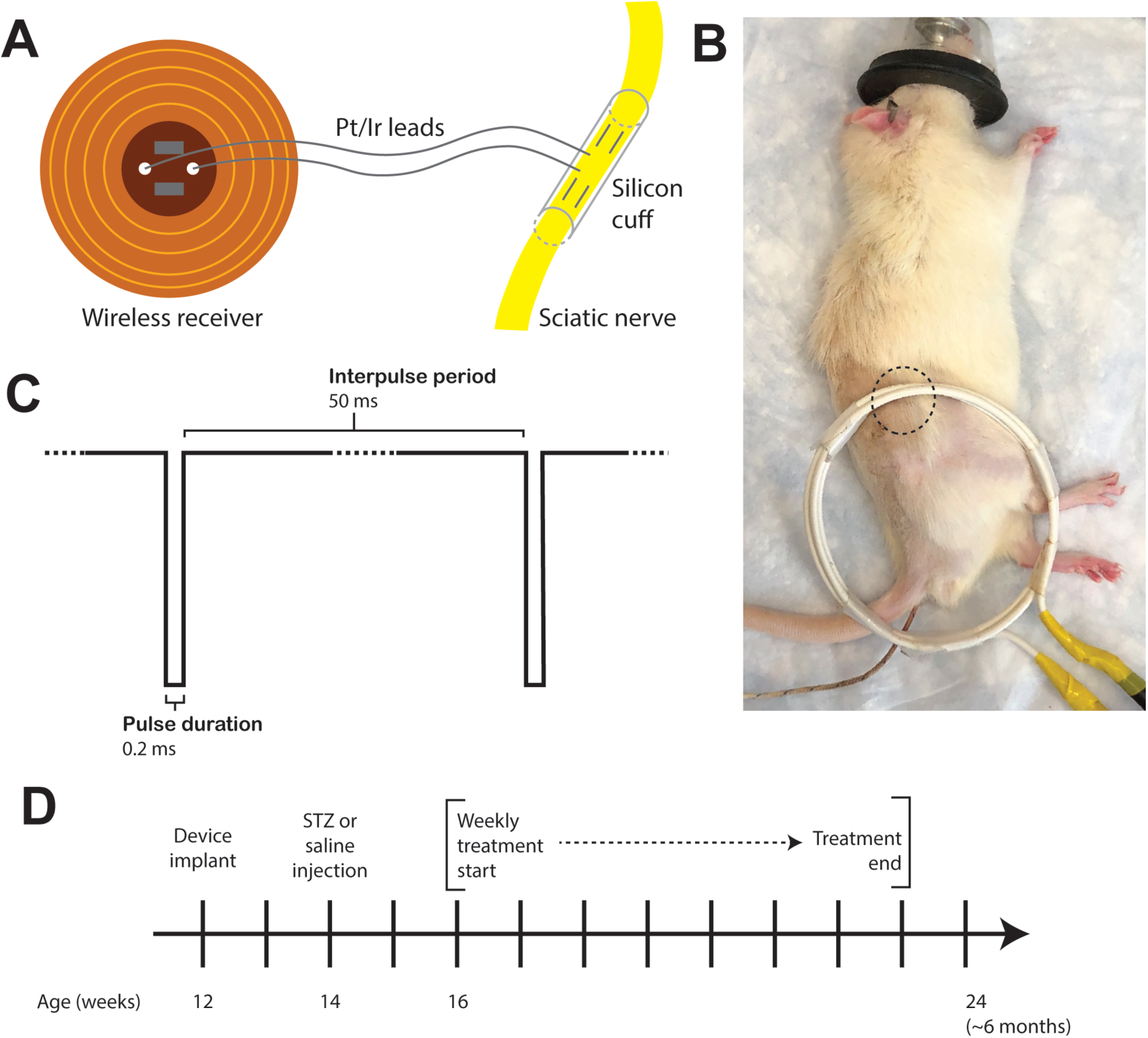
Study design. Fully implantable sciatic nerve cuff stimulators were fabricated by threading braided Pt/Ir leads through 8-mm of silicon tubing (1.5-mm diameter); wires were soldered to a wireless receiver that was then embedded in medical silicon (**A**). The cuff was implanted onto the sciatic nerve and the receiver subcutaneously over the thigh muscles (**B**, dotted circle). The implanted receiver was activated externally by a superimposed transmitter coil while the animals were anesthetized with 2% isoflurane (**B**). The parameters for weekly eStim were one-hour total treatment time with a supramaximal cathodic square pulse stimulus at 20 Hz (**C**). (**D**) Experimental timeline.

### 2.2 Animals

The procedures in this investigation were approved by the Washington University Animal Studies Committee (Saint Louis, MO, USA). Male Lewis rats at 7-weeks of age were obtained from Charles River (Strain 004) and housed on a 12-hour light/dark cycle while fed *ad libitum* (LabDiet, 53WU, PicoLab Rodent Diet 20). To induce T1D, rats at 14-weeks age were fasted for 24-hours on aspen bedding, after which a single 65 mg/kg dose of STZ was administered by IP injection. Controls received saline vehicle. After injection, fasting was continued for 2-hours and rats were given 10% sucrose filter-sterilized water *ad libitum* for 24-hours. After 24-hours, rats were returned to cobb bedding and filtered water. Body mass and tail blood glucose were monitored daily after STZ injection with an electronic scale and with a tail prick by blood glucometer (Bayer Contour Next) for 3-days post-injection. Blood glucose and body mass were subsequently recorded weekly for all animals. Rats exhibiting successive reductions in weight and bradykinesia were given biweekly 2 mL subcutaneous saline injection, wet food and hydrogel until stable body mass was restored. At the end point, rats were euthanized *via* carbon dioxide overdose followed by pneumothorax. Tissues were collected and weighed on an electronic scale. Collected tissues were fixed in 10% neutral buffered formalin (Fisher Scientific 23-245684) for 24-hours prior to processing as detailed below. Tibia length was measured with digital calipers (iKKEGOL).

### 2.3 eStim device implantation and treatment regimen

All surgeries were conducted under anesthesia with 2% isoflurane. All animals were implanted with a device. An incision was made on the lateral surface of the right thigh to expose the sciatic nerve. The silicon cuff was placed around the nerve and closed with one stitch of 6-O nylon suture (McKesson, REF S1698GX), and the receiver was placed subcutaneously proximal to the cuff. Device function was verified at the time of surgery by activating the device with a superimposed transmitter coil (Fig.1B) and the implant site muscle and skin layers were closed with 5-O Vicryl (Ethicon, J303H) and 4-O Nylon (McKesson, REF S662GX) suture, respectively. Animals received a single 1.0 mg/kg dose of buprenorphine sustained-release (ZooPharm) subcutaneously one hour before surgery for post-operative analgesia.

Beginning at 16-weeks of age and continuing for 8-weeks, all animals were anesthetized with 2% isoflurane and those animals assigned to the stimulation group were treated weekly for one hour with a supramaximal cathodic pulse, 0.2 ms pulse duration, and 20 Hz frequency (Fig.1C). These parameters have been shown to enhance neural regrowth post-transection (MacEwan et al., 2018; Geremia et al., 2007). Initially, all animals showed maximal activation of muscles in the lower limb at a threshold of 8-9V at the start of the experiment. Maximal muscular activation effect was defined visually by pointed toes and clenched paw and by palpating twitches in the tibialis anterior and gastrocnemius muscles. Muscular contraction was assessed visually and by palpation during each week of eStim for each animal with a goal to perform stimulations to achieve a maximally-activated effect. Thus, if the effect was not achieved by the initial stimulus intensity, it was increased by raising first the voltage and then the pulse duration in order to achieve the maximally-activated muscle contraction at the start of the experiment.

### 2.4 In vivo computed tomography: cortical bone

All animals were scanned *in vivo* at 24-weeks of age prior to end point. Animals were anesthetized with 1-2% isoflurane and placed into the scanning bed. A piece of VetWrap bandage was taped over the animal’s torso to reduce loss of body heat. The top limb was placed into a rig to straighten and stabilize the leg during the scan with the foot secured. *In vivo* scans were conducted on the mid-diaphysis: a 3-mm region (200 slices) centered halfway between the proximal end of the tibia and the tibiofibular junction (VivaCT40; Scanco Medical; 70 kVp; 114 uA; 15 µm voxel size). After the first limb was scanned, the animal was turned on the bed and the scan process was repeated for the contralateral limb. The total time under anesthesia exceeded no more than one hour. Analysis was performed using the Scanco software. The entire 3-mm ROI at the mid-diaphysis was contoured and analyzed at a threshold of 250 with sigma and support values of 0.8 and 1, respectively. Bone volume fraction (BV/TV), cortical thickness (mm), tissue mineral density (mg HA/cm^3^), total area (mm^2^), bone area (mm^2^), medullary area (mm^2^), and pMOI (mm^4^) were extracted for data analysis.

### 2.5 Single frame motion analysis

Animals were trained to walk across a 3-foot-long, 3.5-inch wide wooden plank to their home cage by placing them on the plank at increasingly distant positions from their home cage. Tickling was used as a reward for task completion (Cloutier et al., 2018). Once animals were adequately trained to walk from the most distant end of the plank to their home cage without pausing, the animals were recorded from behind (iPhone X). Video analysis was performed in MATLAB using a custom guided-user interface designed to measure the first-quadrant angle between the horizontal axis and the foot-to-base vector – this parameter is called the foot-to-base angle (FBA) (Fey et al., 2010). The horizontal axis was determined by the edge of the home cage. The foot-to-base vector was measured by drawing a line between the heel and the midplantar surface at a frame just before pushoff. The angle was measured for every frame available for each foot while the animal was actively walking.

### 2.6 Von Frey

The up-down method was used to assess mechanical allodynia in all animals at 23-weeks of age (Chaplan et al., 1994). Animals were placed on a chicken-wire metal grid fixed to a wooden frame. The frame was elevated 1.5-feet above a countertop to allow for testing and viewing the plantar surface of the paw. A mouse cage was placed over the animal to restrain motion. Animals were allowed to acclimate for 5 minutes, at which point they were no longer actively exploring their environment. Manual von Frey monofilaments ranged from 8-300 grams force. A response was recorded as hindpaw withdrawal upon or subsequent to application of the filament. Filaments were applied alternating sides as described previously, and after a response, animals were allowed to reacclimate for 1 minute.

### 2.7 Ex vivo computed tomography: cancellous bone

Prior to sectioning for histology, explanted tibias were embedded in 2% agarose, and a 6-mm region was scanned starting at the growth plate (VivaCT40; Scanco Medical; 70 kVp; 114 uA; 15 µm voxel size). Analysis was performed using the Scanco software. From 2-mm (133 slices) below the growth plate, an ROI of 1.5-mm (100 slices) was contoured and analyzed at a threshold of 240 with 0.8 sigma and 1 support. Total volume (mm^3^), bone volume (mm^3^), bone volume fraction (BV/TV), structural model index (SMI), connectivity density, trabecular number, trabecular thickness (mm), trabecular separation (mm), and tissue mineral density (mg HA/cm^3^) were extracted for analysis.

### 2.8 Histology and bone marrow adipocyte analysis

All histology was performed by the WUSM Musculoskeletal Histology and Morphometry core. Prior to embedding, tibias were dehydrated in a reverse gradient to 70% ethanol. Tibias were halved at approximately the 50% site between the proximal end and the tibiofibular junction with a rotary saw tool (Dremel). The proximal half of the tibia was fully decalcified in 14% EDTA (Sigma-Aldrich E5134), pH 7.4 prior to paraffin embedding, sectioning (10 µm thickness) and staining with hematoxylin and eosin. Images were taken on a Hamamatsu 2.0-HT NanoZoomer System with NDP.scan 2.5 image software at 20x in brightfield mode.

The H&E stained tibial cross-sectional slides were scanned using a Hamamatsu 2.0-HT NanoZoomer System with NDP.scan 2.5 image software. The acquired images were exported as TIFF files under 10x magnification and were processed in Fiji to measure average adipocyte cell size and number. Briefly, the scale in Fiji was first set to be consistent with the original image (1.084 pixels/µm). The image was then converted to 8-bit and the cortical bone was specifically selected by thresholding. A median filter with a radius of 10 pixels was applied. The image was then inverted and the area of bone marrow cavity was selected and measured using the ‘Wand tool’ and the ‘Measure’ command. A threshold of 230 to 255 was applied to the original 8-bit image and everything outside the bone marrow cavity was cleared using the ‘Clear Outside’ command. A median filter with a radius of 2 pixels was applied to the image and the non-adipocyte structures were selected and eliminated using the ‘Analyze Particles’ tool by setting the circularity to 0 – 0.2. The cleaned image was further processed using the ‘Watershed’ tool and the adipocyte size and number were finally determined using the ‘Analyze Particles’ tool by setting the size to 200 to 4000 μm^2^ and circularity to 0.50 – 1.00. The average adipocyte cell size and number per bone marrow area were calculated using Excel.

### 2.9 Statistics

Statistical analyses for this study were performed in GraphPad Prism. Tests included two-way ANOVA, three-way ANOVA, and mixed effects analyses. Specific information on statistical tests is detailed in the figure legends. A p-value of less than 0.050 was considered statistically significant. Quantitative assessments including bone length, organ mass, microcomputed tomography, behavioral assessments, and bone marrow adiposity measurements were performed by individuals blinded to the experimental groups.

## 3 Results

### 3.1 eStim device design and experimental timeline

Fully implantable sciatic nerve cuff stimulators were fabricated by threading braided Pt/Ir leads through 8-mm of silicon tubing (1.5-mm diameter); leads were soldered to a wireless receiver (Fig.1A; (Gamble et al., 2016). As the average diameter of the rat sciatic nerve is approximately 1-mm (Isaacs et al., 2014; Onode et al., 2019), it was neither expected nor intended that the cuff would constrict the sciatic nerve at any stage of development. The cuff was implanted unilaterally onto the right sciatic nerve and the receiver was placed subcutaneously over the right thigh muscles (Fig.1B, dotted circle). All animals received a cuff implant at 12-weeks of age. Diabetes was induced at 14-weeks of age by a single 65 mg/kg intraperitoneal injection of streptozotocin (STZ). In the eStim groups, the implanted receiver was activated externally by a superimposed transmitter coil (Fig.1B). Weekly, one-hour eStim treatment consisted of a supramaximal cathodic square pulse stimulus at 20 Hz (Fig.1C), as defined previously for nerve regeneration (MacEwan et al., 2018; Geremia et al., 2007). Weekly eStim or sham (anesthesia-only) treatments began 2-weeks post-STZ and continued for 8-weeks prior to end point analysis at 24-weeks of age (Fig.1D). There are four groups in this study: control sham, control eStim, T1D sham, and T1D eStim.

### 3.2 Regulation of blood glucose, body and tissue mass by T1D and eStim

Blood glucose and body mass were monitored longitudinally to confirm diabetic induction and sustained hyperglycemia throughout the study. In T1D rats, blood glucose increased from 106±7 mg/dL to 503±70 mg/dL post-injection with STZ (Fig.2A). Sustained hyperglycemia was maintained throughout the course of the experiment (Fig.2A). By contrast, control animals remained normoglycemic (Fig.2A). From the time of T1D onset at 14-weeks of age to the final week of treatment, body mass increased by 28±4% in control animals and decreased by 23±4% in diabetic animals relative to baseline (Fig.2B). As expected, unilateral sciatic nerve eStim for 1-hour per week did not influence body mass or blood glucose in healthy or diabetic animals, relative to sham, anesthesia only controls (Fig.2A-B).

**Figure 2.**
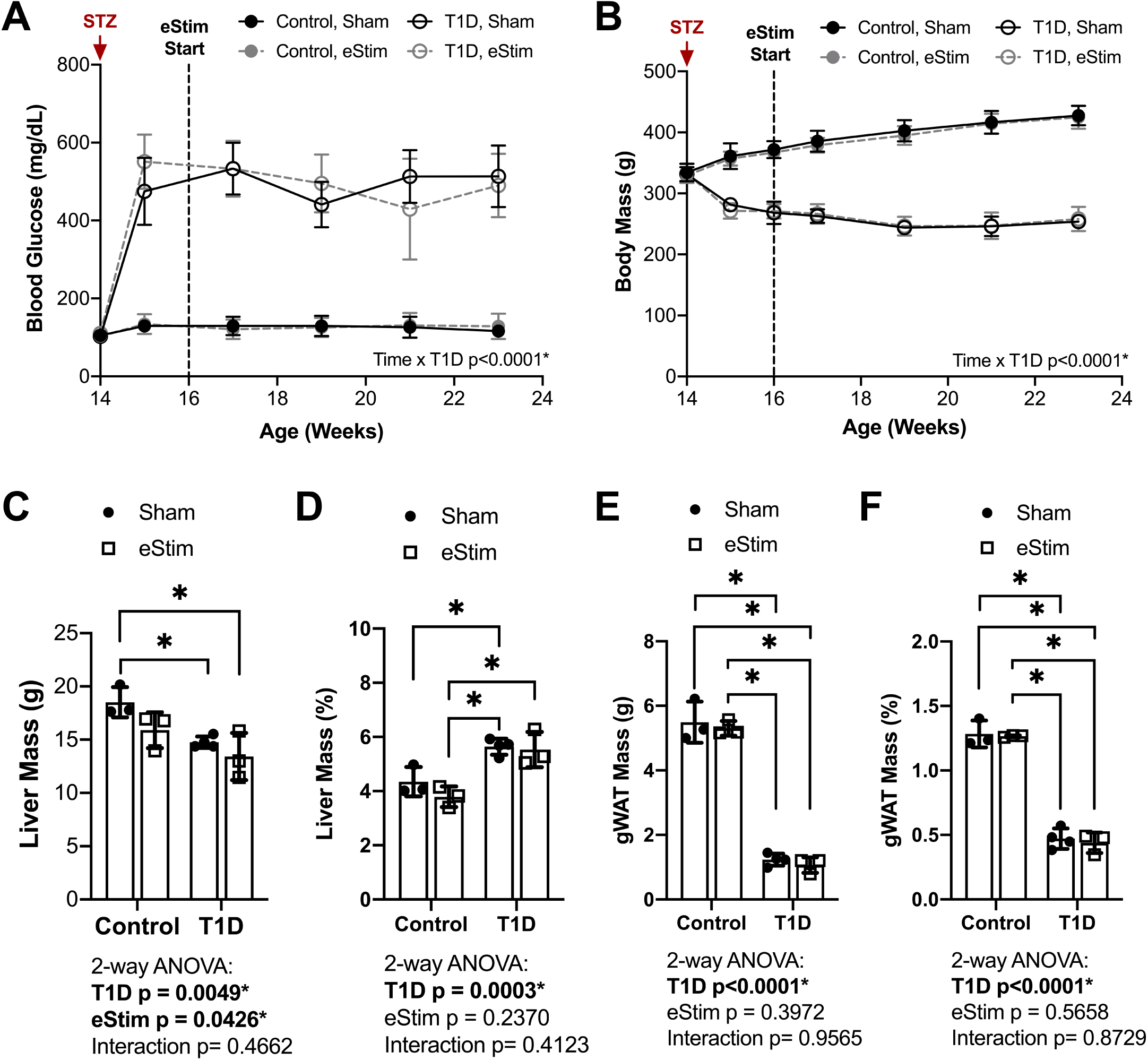
Blood glucose and body and organ mass. Blood glucose and body mass were evaluated starting at the time of STZ induction (14-weeks of age) up to the final week of treatment (24-weeks of age). (**A**) Blood glucose. (**B**) Body mass. (**C**) Liver mass at end point. (**D**)Liver mass normalized to body mass. (**E**) Gonadal white adipose tissue (gWAT) mass at end point. (**F**) gWAT mass normalized to body mass. Statistics for blood glucose and body mass were performed by mixed effects and three-way ANOVA analyses, respectively, as a single missing value was present in the blood glucose data; Control, sham n = 7; Control, eStim n = 6; Diabetic, sham n = 8; Diabetic, eStim n = 5; *p<0.050. Statistics for organ masses was performed by two-way ANOVA with Sidak’s multiple comparisons test; Control, sham n = 3; Control, eStim n = 3; Diabetic, sham n = 4; Diabetic, eStim n = 3; *p<0.050.

At the end point, tissues including liver, gonadal white adipose tissue (gWAT), and spleen were dissected and weighed to gauge overall health. T1D resulted in reduced absolute liver and gWAT mass (Fig.2C,E). When normalized to body weight, liver mass was elevated by 37% in T1D rats (Fig.2D) and the relative quantity of gWAT was reduced by 78% (Fig.2F). Bioelectric nerve stimulation resulted in reduced absolute liver mass, but normalization to body weight eliminated this effect (Fig.2C-D). Bioelectric nerve stimulation did not alter gWAT (Fig.2E-F). T1D caused absolute decreases in spleen mass that were proportional to body size and not impacted by eStim (data not shown). Overall, this confirmed that intermittent, unilateral stimulation of the sciatic nerve did not cause overt global changes in peripheral tissues.

Stimulation of the sciatic nerve causes unilateral muscle contraction (Supplemental Video S1). We hypothesized that this may be sufficient to increase the muscle mass in healthy animals and to rescue muscle atrophy in those with T1D. In previous studies, type II (fast-twitch) muscle fibers have displayed distinct effects compared to type I (slow-twitch) fibers in the STZ-induced model of T1D (Rutschmann et al., 1984; Cotter et al., 1989). Thus, muscles of predominately type I (soleus (SOL)), type II (tibialis anterior (TA), extensor digitorum longus (EDL), plantaris (PL)) and mixed type fiber compositions (lateral and medial gastrocnemius (LGC/MGC)) were selected for analysis (Fig.3A). Rats with T1D had reduced hindlimb muscle mass relative to controls (Fig.3B-F). Specifically, in diabetic animals, muscle masses were bilaterally reduced by 32% in the soleus (Fig.3B); by 56%, 60%, and 54% in the tibialis anterior, EDL and plantaris, respectively (Fig.3C-E); and by 53% in the gastrocnemius (Fig.3F). Consistent with previous reports (Cotter et al., 1989; Rutschmann et al., 1984), diabetic muscles containing a significant population of type II fibers were more severely affected than those with predominately type I fibers. In addition to the effect of diabetes, we also considered the effects of sciatic nerve cuff placement and eStim using 3-way ANOVA (T1D x Cuff x eStim). The placement of a sciatic nerve cuff did not independently influence muscle mass. In addition, contrary to our expectations, eStim treatment did not significantly alter or improve muscle mass in control rats or those with T1D (Fig.3B-F).

**Figure 3.**
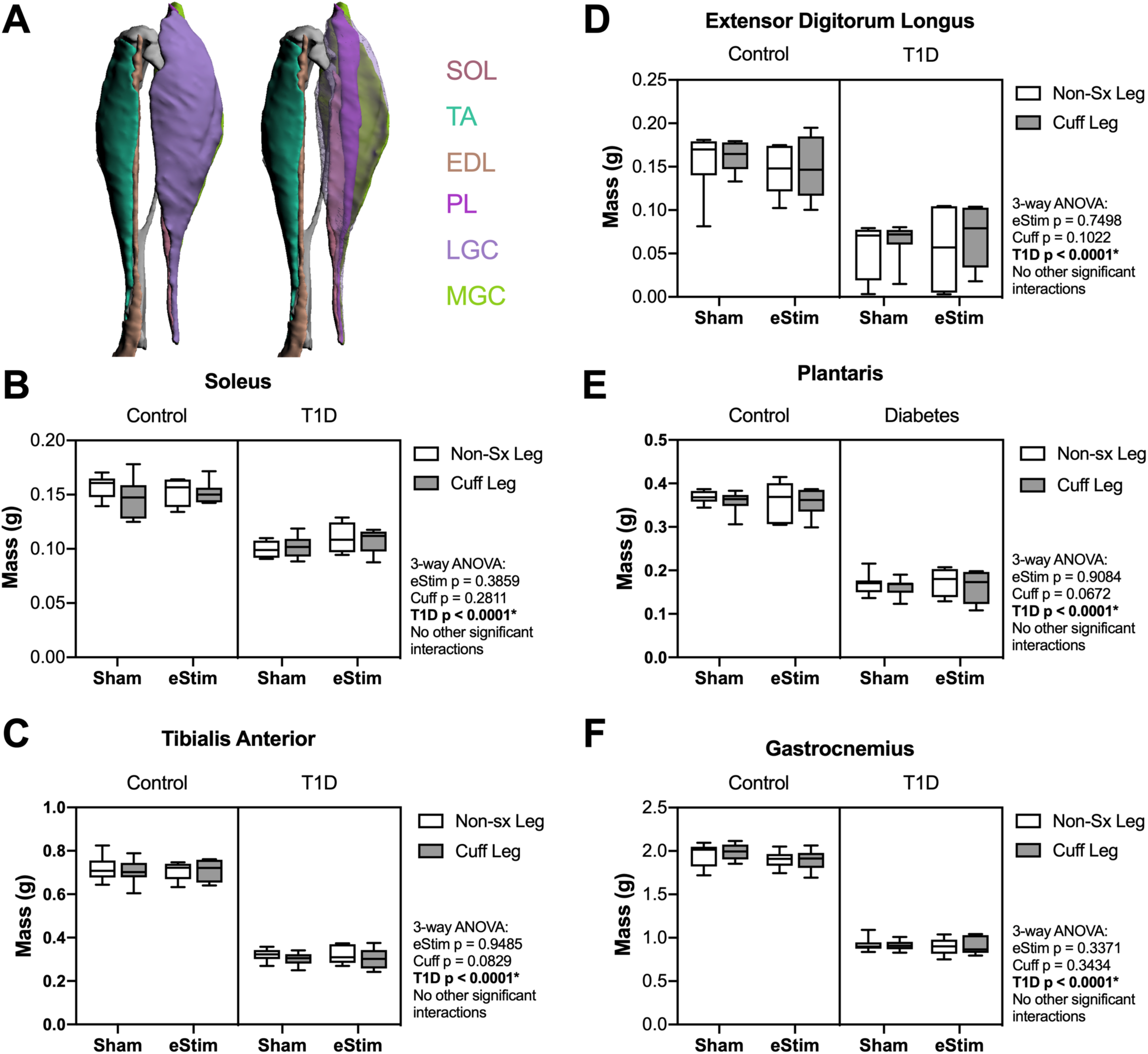
Muscle mass. At end point, muscles surrounding the tibia (**A**) were dissected and weighed. (**B**) Soleus (SOL) muscle mass. (**C**) Tibialis anterior (TA) muscle mass. (**D**) Lateral and medial gastrocnemius (LGC, MGC) combined muscle mass. (**E**) Extensor digitorum longus (EDL) muscle mass. (**F**) Plantaris (PL) muscle mass. Control, sham n = 7; Control, eStim n = 6; Diabetic, sham n = 8; Diabetic, eStim n = 5; Three-way ANOVA; *p<0.050. Panel A generated from (Charles et al., 2016).

### 3.3 Gait alterations and mechanical allodynia with T1D, sciatic nerve cuff, and eStim

As the sciatic nerve is involved in locomotion, its manipulation could result in gait changes. For example, foot-to-base angle (FBA) is reduced in rodents with sciatic nerve damage (Fey et al., 2010). To assess this, single-frame motion analysis was conducted at 11-weeks (baseline) and 24-weeks of age (end point). FBA was measured as the first-quadrant angle between the line from midplantar surface to heel and the perpendicular (home cage edge) (Fig.4A; (Kruspe et al., 2014; Fey et al., 2010). At baseline, prior to surgery, FBA was not significantly different between the left and right limbs (Fig.4B). At the end point, T1D resulted in a bilateral reduction of FBA by -12%, independent of cuff placement or eStim (Fig.4C). FBA was further reduced unilaterally on the cuffed side of non-stimulated animals by -12±10% in controls and by -13±14% in diabetics (Fig.4C). In stimulated animals, however, this effect was suppressed. The FBA of the cuffed limb was partially restored in control and diabetic eStim groups to -3±7% and -3±13%, respectively, relative to the control side (Fig.4C). Overall, these results show a unilateral, detrimental effect of sciatic nerve cuffing on gait that is partially restored by eStim.

**Figure 4.**
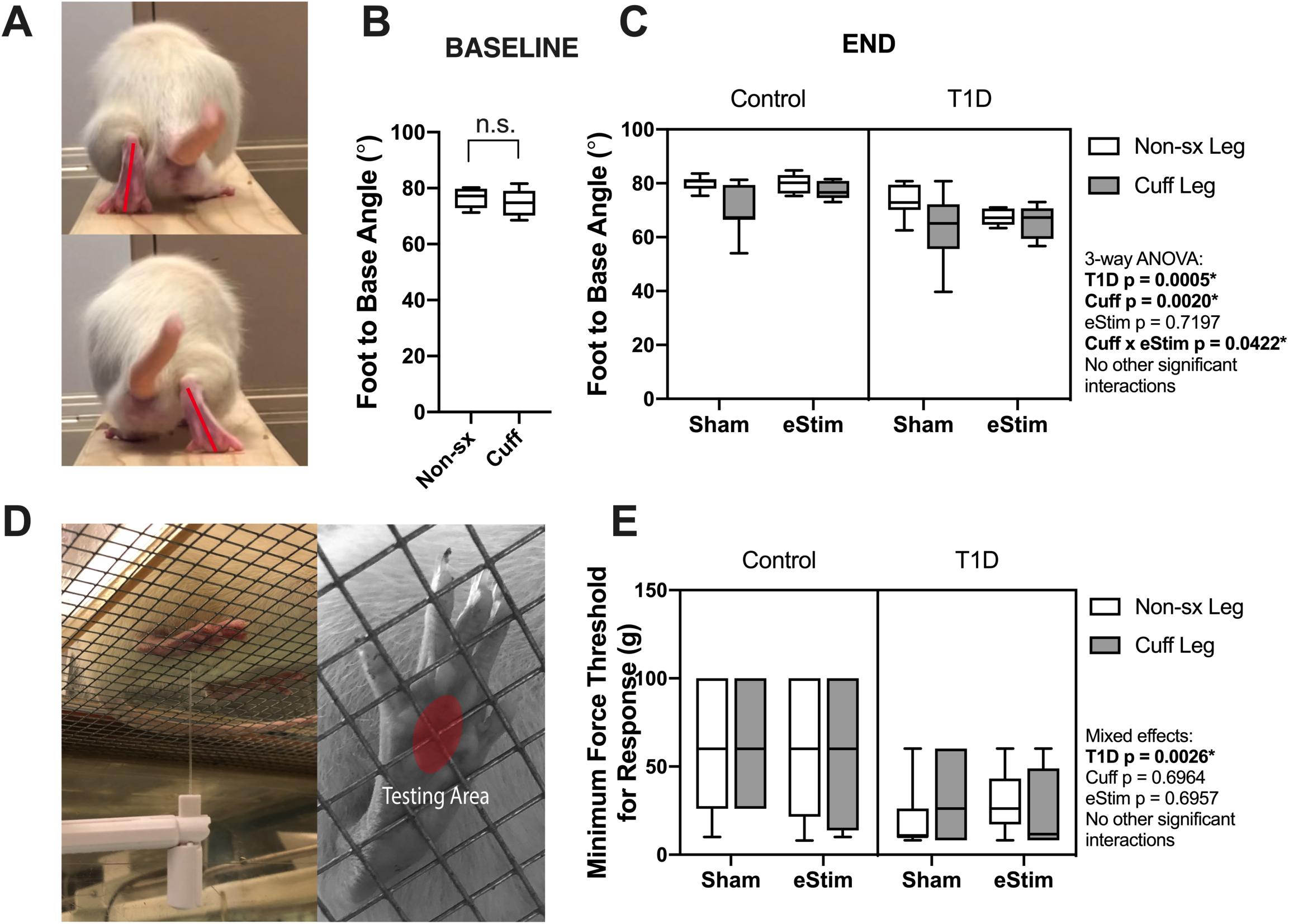
Neuromuscular behavioral assessments. (**A**) Foot to base angle (FBA) was measured using single frame motion analysis (SFMA). (**B**) Baseline FBA measurements. (**C**) End point FBA measurements. Manual von Frey filaments were also applied to the midplantar surface of the hindpaw (indicated by a red circle) (**D**). (**E**) Minimum force threshold for response at end point. Statistics for baseline FBA measurements was performed by paired t-test. Control, sham n = 7; Control, eStim n = 6; Diabetic, sham n = 8; Diabetic, eStim n = 5; Three-way ANOVA; *p<0.050.

Rodent models of T1D exhibit mechanical allodynia, representative of sensory DPN (Morrow, 2004). We hypothesized that application of neuroregenerative eStim would oppose the progression of DPN, resulting in normalization of mechanical sensitivity. Cuff placement can also independently cause unilateral sensitivity if placed too tightly around the nerve (Austin et al., 2012; Mosconi and Kruger, 1996). Thus, we measured mechanical allodynia at 23-weeks of age (one week before end point) using manual von Frey filaments applied to the mid-plantar surface of the hindpaw (Fig.4D, red oval; (Chaplan et al., 1994). As expected, diabetic animals exhibited, on average, a 56% reduction in the response threshold bilaterally compared to controls, indicative of increased sensitivity to mechanical stimuli (Fig.4E, 3-way ANOVA, T1D p=0.0026). However, contrary to our expectations, eStim treatment did not alter the mechanical sensitivity of the controls or rescue the mechanical allodynia of those with T1D (Fig.4E, 3-way ANOVA, eStim p=0.6957). The presence of the unilateral sciatic nerve cuff did not impact the mechanical sensitivity in the cuffed limb in either control or diabetic animals, confirming the absence of overt nerve constriction or irritation in our model (3-way ANOVA, Cuff p=0.6964).

### 3.4 Changes in bone and bone marrow adiposity with T1D, sciatic nerve cuff and eStim

#### 3.4.1 Bone length

Sensory neurotransmitters such as calcitonin gene related peptide (CGRP) have previously been shown to promote bone formation in developing animals (Xu et al., 2020). By contrast, T1D can limit bone growth (Silva et al., 2009). To assess the effects of T1D, cuff placement, and eStim on skeletal growth, left and right side tibial lengths were measured with digital calipers and compared using 3-way ANOVA (T1D x Cuff x eStim). Tibial length was decreased bilaterally by 5% in rats with T1D when compared to controls, independent of cuff placement or eStim (Fig.5A, T1D p<0.0001). In addition, the cuffed limb was slightly shorter than the non-surgical limb in sham treated controls (−0.3% and -0.5% in healthy control and diabetic animals, respectively; Fig.5A). By contrast, in animals treated with unilateral chronic eStim the cuffed limb was slightly longer (+0.3% and +1.6% in control and diabetic animals, respectively) (Fig.5A), indicating an interaction between sciatic nerve cuffing and eStim on longitudinal bone growth (3-way ANOVA, Cuff x eStim p=0.0243).

**Figure 5.**
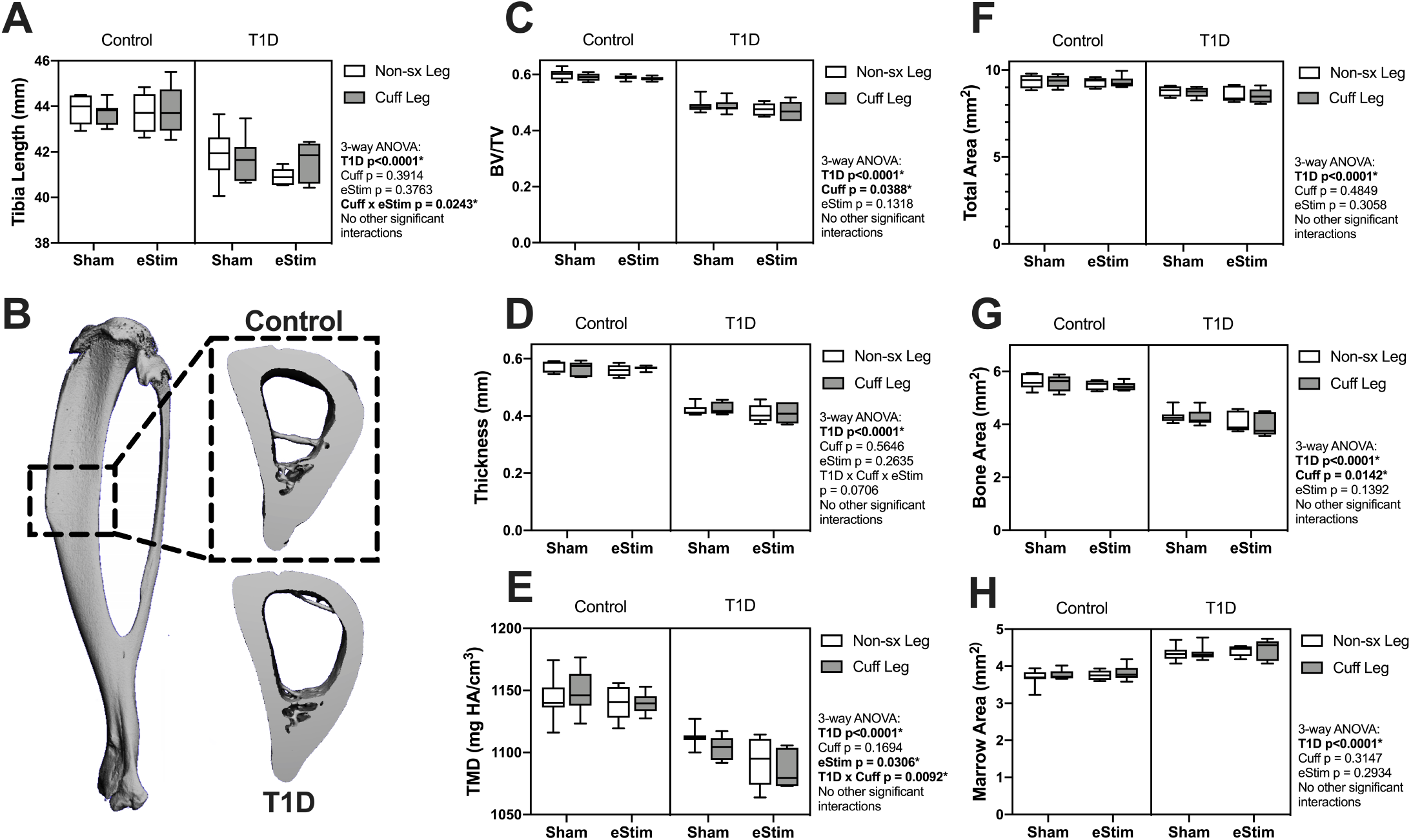
Tibia length and cortical bone. (**A**) End point tibial length as assessed after dissection using digital calipers. Cortical bone was evaluated by µCT in a 3-mm region centered midway between the proximal end of the tibia and the tibia-fibula junction (**B**). (**C**) Cortical bone volume fraction (BV/TV). (**D**) Cortical thickness. (**E**) Cortical tissue mineral density (TMD). (**F**) Cortical total area. (**G**) Cortical bone area. (**H**) Cortical marrow area. Control, sham n = 7; Control, eStim n = 6; Diabetic, sham n = 8; Diabetic, eStim n = 5; Three-way ANOVA; *p<0.050.

#### 3.4.2 Cortical bone

As mentioned above, sensory nerves in bone are thought to release anabolic neuropeptides near bone-forming cells (Brazill et al., 2019; Tomlinson et al., 2016); thus, we expected that DPN prevention and/or neural activation by eStim would increase bone mass. Cortical bone was analyzed by *in vivo* microcomputed tomography at 12-weeks (baseline) and 24-weeks of age (end point) in a 3-mm section of the mid-diaphysis, centered between the proximal end of the tibia and the tibia-fibula junction (Fig.5B). At baseline, prior to cuff implantation, the left and right limbs exhibited no difference in bone size or morphology (data not shown). Cortical tissue mineral density (TMD) was 0.8% lower in the right limb (the limb to be cuffed) relative to the left at baseline (p=0.0412).

At end point, rats with T1D demonstrated a 19% reduction in cortical bone volume fraction (BVF), a 26% decrease in cortical thickness, and a 4% decrease in cortical TMD relative to non-diabetic controls (Fig.5C-E). Reduced bone quantity in diabetic animals was driven by a 24% reduction in bone area, a 7% decrease in total area, and a 16% increase in marrow area (Fig.5F-H). Sciatic nerve cuffing over 12-weeks had a small negative effect on cortical BVF in the cuffed limb (ranging from -0.4 to -1.5%), independent of eStim or T1D (Fig.5C). This effect was also reflected by a unilateral 1 to 3% reduction in bone area in the cuffed limb relative to the non-surgical control side (Fig.5G). The only observed effect of eStim on cortical bone was a 0.6% and 1.6% bilateral reduction in TMD in control and diabetic animals, respectively, relative to non-stimulated, sham controls (Fig.5E). In summary, T1D resulted in reduced cortical bone quantity and mineral density that was not rescued by eStim. In fact, eStim and sciatic nerve cuffing introduced additional cortical bone deficits in both control and diabetic animals including bilaterally reduced TMD (as a result of eStim) and unilaterally reduced bone mass (as a result of sciatic nerve cuffing).

#### 3.4.3 Cancellous bone

The inside of the long bones is filled with spongy, cancellous bone that is concentrated largely at the metaphyses. This bone has a high turnover rate and is susceptible to systemic change, including well-documented decreases in rodents with T1D (Silva et al., 2009). To assess the effects of T1D, cuff placement and eStim on metaphyseal cancellous bone, we analyzed a 1.5-mm region starting 2-mm below the growth plate (Fig.6A). Consistent with previous reports, cancellous BVF was reduced by 31% in rats with T1D (Fig.6B). This was associated with a 10% increase in trabecular number and a 25% decrease in trabecular thickness (Fig.6C-D). Animals with T1D exhibited a trending 20% reduction in connectivity density (data not shown; 3-way ANOVA, p=0.0651) and a significant increase in structure model index (SMI) (Fig.7E; 2.6 vs. 3.1; SMI = 0 for plates, 3 for rods and 4 for solid spheres). Cancellous bone mineral density (BMD) was not significantly different between groups (data not shown). Unlike the effects observed with T1D, cancellous bone quantity, morphology, and mineralization were not modified by sciatic nerve cuffing or chronic eStim.

**Figure 6.**
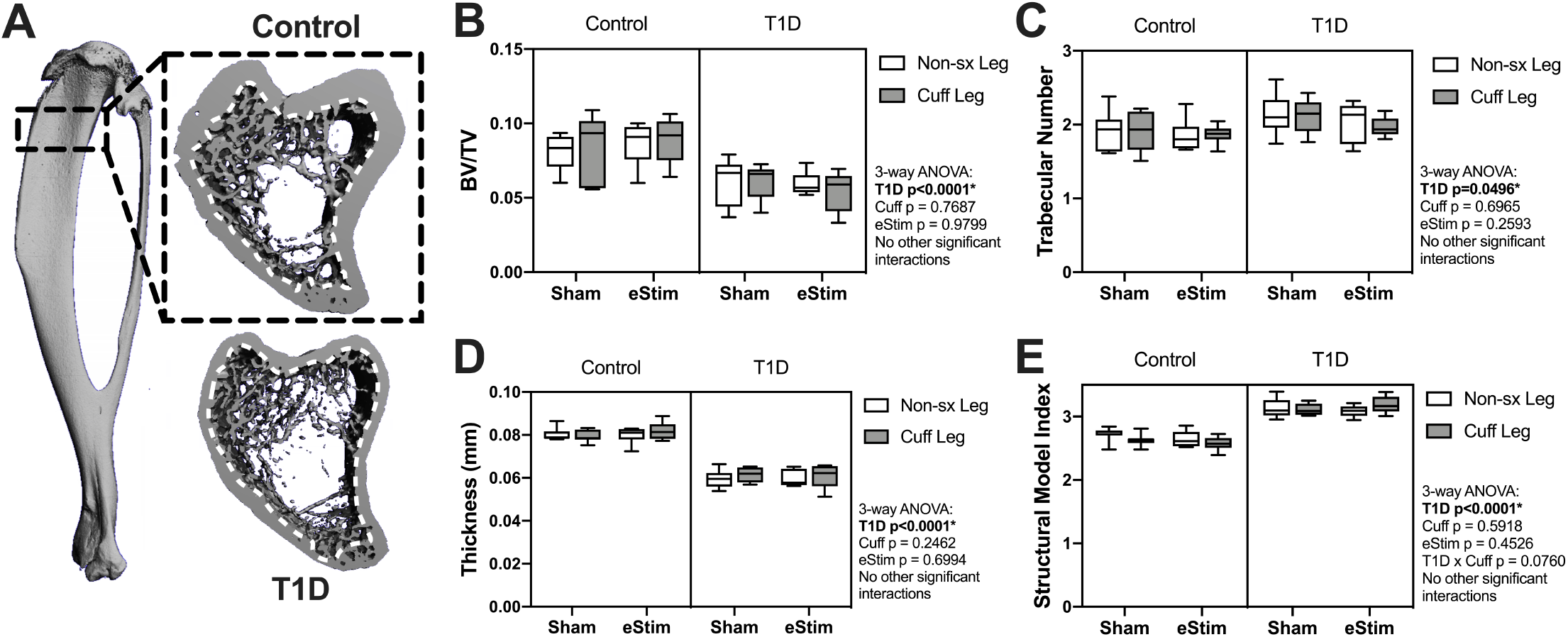
Cancellous bone. Cancellous bone was analyzed 2-mm below the growth plate in a 1.5-mm region (**A**), dotted line represents contour inside of which bone was analyzed). (**B**) Cancellous bone volume fraction (BV/TV). (**C**) Trabecular number. (**D**) Trabecular thickness. (**E)**Structural model index (SMI). Control, sham n = 7; Control, eStim n = 6; Diabetic, sham n = 8; Diabetic, eStim n = 5; Three-way ANOVA; *p<0.050

**Figure 7.**
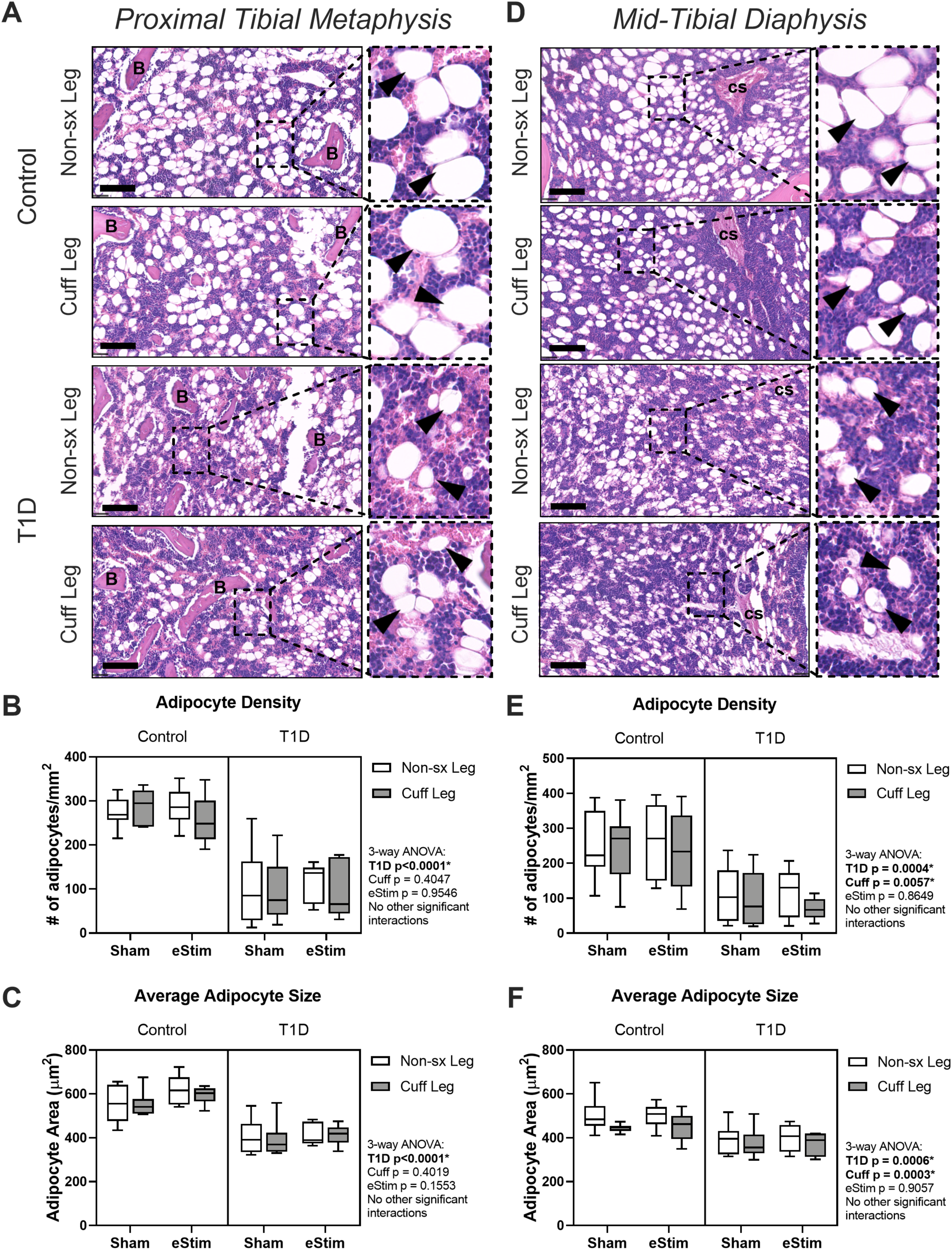
Bone marrow adipose tissue. Bone marrow adiposity was analyzed in the metaphysis 2-mm below the growth plate and in the diaphysis 2-mm above the tibial crest. (**A**) BMAT histology in the metaphysis in healthy and control animals. (**B**) Adipocyte density in the metaphysis. (**C**) Average adipocyte size in the metaphysis. (**D**) BMAT histology in the diaphysis in health and control animals. (**E**) Adipocyte density in the diaphysis. (**F**) Average adipocyte size in the diaphysis. B=bone; cs = central venous sinus. Scale = 100 µm. (Control, sham n = 7; Control, eStim n = 6; Diabetic, sham n = 8; Diabetic, eStim n = 5) Three-way ANOVA; *p<0.050.

#### 3.4.4 Bone marrow adiposity

In addition to bone, the skeleton is filled with a unique population of adipocytes that are collectively known as the bone marrow adipose tissue (BMAT). BMAT is an emerging regulator of hematopoietic, metabolic and skeletal health (Scheller et al., 2016). The ability of nerve stimulation to regulate bone marrow adipocytes *in vivo* remains unknown. In this study, bone marrow adipocytes were analyzed in the metaphysis approximately 2-mm below the growth plate (Fig.7A-C) and in the diaphysis approximately 2 mm above the peak of the tibial crest (Fig.7D-F), matching our regions of skeletal morphologic assessment in Fig.6 and Fig.5, respectively. T1D and sciatic nerve cuff placement both had independent effects on BMAT (Fig.7); however, the effect of sciatic nerve cuffing was region-specific. In the metaphysis, T1D resulted in a 59% reduction in BMA number, a 63% decrease in BMA density, and a 30% reduction in BMA size (Fig.7A-C and data not shown). Neither sciatic nerve cuffing nor eStim resulted in a change in metaphyseal BMAT. In the diaphysis, T1D similarly resulted in a 39% reduction in BMA number, a 59% decrease in BMA density, and a 19% reduction in BMA size (Fig.7D-F and data not shown). However, contrary to the metaphyseal data, sciatic nerve cuffing alone also caused a 5-32% reduction in BMA number, a 5-38% reduction in BMA density, and a 4-12% reduction in adipocyte cell size relative to the contralateral limb (Fig.7D-F and data not shown). These cuff-side effects on bone marrow adiposity were driven primarily by changes in the non-diabetic control group and align spatially with previously reported differences in the innervation of the bone marrow adipocyte population along the length of the limb (Wee et al., 2019). Bioelectric stimulation did not modify the effects of T1D or cuff placement on bone marrow adiposity.

## 4 Discussion

Despite promising results showing the pro-regenerative benefits of eStim (Geremia et al., 2007; Gamble et al., 2016; Wong et al., 2015), studies to date have not addressed its utility to prevent the onset of neuropathic symptoms and musculoskeletal decline in the context of metabolic diseases like T1D. Our experimental design allowed us to address the above questions using a fully implantable, wireless sciatic nerve stimulation device in a rat model of T1D while also considering the effects of cuff placement and chronic eStim in otherwise healthy animals.

Rodent models of T1D are characterized by progressive increases in mechanical sensitivity in addition to deficits in bone quantity, muscle mass and neuromuscular function (Cotter et al., 1989; Morrow, 2004; Silva et al., 2009). Similar to previous studies, we observed a significant decrease in hindlimb muscle mass, reduced withdrawal threshold to mechanical stimuli in the hindpaw, changes in gait, and substantial decreases in bone in animals with T1D. These results are entirely consistent with previous studies with one exception. Previous reports of STZ-induced T1D in rodents have demonstrated an increase in bone marrow adiposity and a decrease in peripheral fat mass (Botolin et al., 2005; Martin and McCabe, 2007). While we did observe a significant reduction in peripheral fat, we did not observe an increase in tibial bone marrow adipocyte density or size in either the metaphysis or diaphysis of rats after 10-weeks of STZ-induced T1D. In fact, contrary to prior results, both adipocyte density and size were substantially reduced in diabetic rats. Previous findings were observed in mice at 4- to 6-weeks post-induction; thus, our results may indicate a differential effect on bone marrow adiposity with sustained T1D that warrants further investigation.

In our study we also assessed the effect of chronic placement of a polyethylene sciatic nerve cuff for 12-weeks. We found that cuff placement contributed to independent changes in gait, tibial length, cortical bone and bone marrow adiposity. Cuff placement can be used as model of chronic constriction injury (CCI) and neuropathic pain, but the largest inner diameter used in CCI models is approximately half of that used in our device (0.86 vs. 1.5 mm) (Austin et al., 2012; Balasubramanyan et al., 2006; Mosconi and Kruger, 1996). In addition, we did not see a unilateral increase in mechanical sensitivity induced by the placement of a sciatic nerve cuff, a typical indicator of neuropathy in CCI studies. Thus, it is unlikely that the effects on gait and bone were caused by overt nerve constriction. Instead, we anticipate that these results reflect local changes in nerve swelling, fibrosis and inflammation that occur after implantation of a local neuroprosthesis, contributing to the observed changes in gait and unilateral reductions in tibia length, cortical bone mass, and bone marrow adiposity. More sophisticated designs for chronic neural implants, including helical or spiral cuff electrodes, have demonstrated improved flexibility and reduced activation of inflammatory responses (Christie et al., 2017; Günter et al., 2019). Additional research is needed to investigate whether these improved designs limit the undesirable effects on bone as observed in this study.

Beyond consideration of the effects of T1D and cuff placement, our primary goal of this study was to isolate the impact of chronic, intermittent eStim on nerve and bone health and on the progression of DPN. Contrary to our initial hypotheses, we did not observe an effect of eStim on muscle mass, bone quantity, or mechanical sensitivity in healthy animals or a rescue in those with T1D. However, we did find that treatment with eStim opposed unilateral cuff-induced effects on tibial length and gait. This suggests that intermittent application of a local bioelectric stimulus can counteract some of the negative side-effects of cuff placement. Local, stimulation-induced release of anabolic neuropeptides at the growth plate may be involved in rebalancing tibial length (Brazill et al., 2019). Likewise, bioelectric nerve stimulation has been shown to drive changes in muscular gene expression (Brownson et al., 1988), which may underlie functional changes observed in gait independent of muscle mass. Future investigation is required to determine if eStim modulates endochondral ossification at the skeletal growth plate and aspects of muscle function, including underlying gene expression. In addition to these positive effects, unexpectedly, treatment with eStim resulted in a 1-2% bilateral reduction in cortical bone tissue mineral density, independent of T1D. The mechanism underlying this effect remains unknown. However, we hypothesize that nerve action potentials produced by the device used here may have travelled bidirectionally along the axon, resulting in bilateral reductions in skeletal mineralization through activation of central neural relays. Additional studies may address whether this effect was centrally mediated by eStim responses in the spinal cord or brain, or by treatment-induced systemic stress factors otherwise insufficient to alter body mass or blood glucose.

### 4.1 Limitations

The eStim paradigm employed here was previously optimized to enhance neuroregeneration in a post-injury setting (MacEwan et al., 2018; Geremia et al., 2007; Singh et al., 2015), but it was unknown if this paradigm could be leveraged to prevent neuropathy or other musculoskeletal deficits associated with chronic metabolic disease. While it rescued cuff-induced deficits in tibial length and gait, eStim did not prevent T1D-associated mechanical allodynia, osteopenia or sarcopenia. To the extent that our assays could measure, we did not observe a benefit of eStim therapy for nerve function in T1D animals. However, our interpretation is limited by assay specificity. Von Frey and gait analyses are indicative of large diameter sensory and/or motor fiber dysfunction. While our analyses show that eStim did not prevent large fiber neuropathy, it is left to future investigation to determine whether eStim can protect against small-fiber neuropathy. Our study was also limited to a discrete selection of eStim parameters; it remains possible that optimization of these parameters for selective recruitment of large or small caliber axons could better target muscle or bone for therapeutic purposes. A common application of eStim clinically is to treat peripheral pain, in which parameters are typically optimized to block aberrant nerve activity, at either the peripheral or spinal level (Mobbs et al., 2007; Kumar et al., 2008). As such, it would be worthwhile to assess bone health in pre-clinical and clinical models that block neuronal activation, in addition to those designed for activation.

### 4.2 Conclusions

Overall, the stimulation parameters and treatment regimen selected for this study were insufficient to prevent T1D-induced osteopenia, sarcopenia and neuropathy. However, our results indicate that cuff device placement on peripheral nerves can unilaterally reduce cortical bone mass and regulate bone marrow adiposity. In addition, while bioelectric nerve stimulation restored tibial length and gait imbalances induced by sciatic nerve cuffing, it also caused bilateral reductions in cortical bone mineral density. Altogether, this suggests that skeletal health should be monitored in long-term clinical applications of neuromodulation devices to prevent unintended consequences including decreased bone mineral density and increased fracture risk.

## 6 Author Contributions

ATB – Conceptualization, Data curation, Formal analysis, Investigation, Methodology, Project administration, Visualization, Validation, Writing, Reviewing, Editing

IS – Data curation, Formal analysis, Investigation, Visualization, Reviewing, Editing

XZ – Data curation, Formal analysis, Investigation, Methodology, Validation, Visualization, Reviewing, Editing

KM – Data curation, Project administration, Investigation, Supervision, Reviewing, Editing YY – Investigation, Methodology, Resources, Supervision, Reviewing, Editing

MM – Conceptualization, Methodology, Resources, Reviewing, Editing

ELS – Conceptualization, Formal analysis, Funding acquisition, Methodology, Project administration, Resources, Supervision, Visualization, Writing, Reviewing, Editing

## 8 Acknowledgements

We would like to give special thanks to Priscilla Stecher for her help executing treatments in initial pilot studies and Nathan Birenbaum for his practical guidance in eStim device fabrication. We would also like to thank Dr. Matt Ward for providing practical and theoretical training on electrical peripheral nerve stimulation and Dr. Gretchen Meyer for training and assistance in dissecting hindlimb muscles.

## References

Austin, P. J., Wu, A., and Moalem-Taylor, G. (2012). Chronic constriction of the sciatic nerve and pain hypersensitivity testing in rats. J. Vis. Exp. doi:10.3791/3393.

Balasubramanyan, S., Stemkowski, P. L., Stebbing, M. J., and Smith, P. A. (2006). Sciatic chronic constriction injury produces cell-type-specific changes in the electrophysiological properties of rat substantia gelatinosa neurons. J. Neurophysiol. 96, 579–590. doi:10.1152/jn.00087.2006.

Beeve, A. T., Brazill, J. M., and Scheller, E. L. (2019). Peripheral neuropathy as a component of skeletal disease in diabetes. Curr. Osteoporos. Rep. 17, 256–269. doi:10.1007/s11914-019-00528-8.

Botolin, S., Faugere, M.-C., Malluche, H., Orth, M., Meyer, R., and McCabe, L. R. (2005). Increased bone adiposity and peroxisomal proliferator-activated receptor-gamma2 expression in type I diabetic mice. Endocrinology 146, 3622–3631. doi:10.1210/en.2004-1677.

Brazill, J. M., Beeve, A. T., Craft, C. S., Ivanusic, J. J., and Scheller, E. L. (2019). Nerves in bone: evolving concepts in pain and anabolism. J. Bone Miner. Res. 34, 1393–1406. doi:10.1002/jbmr.3822.

Brownson, C., Isenberg, H., Brown, W., Salmons, S., and Edwards, Y. (1988). Changes in skeletal muscle gene transcription induced by chronic stimulation. Muscle Nerve 11, 1183–1189. doi:10.1002/mus.880111113.

Brushart, T. M., Jari, R., Verge, V., Rohde, C., and Gordon, T. (2005). Electrical stimulation restores the specificity of sensory axon regeneration. Exp. Neurol. 194, 221–229. doi:10.1016/j.expneurol.2005.02.007.

Chaplan, S. R., Bach, F. W., Pogrel, J. W., Chung, J. M., and Yaksh, T. L. (1994). Quantitative assessment of tactile allodynia in the rat paw. J. Neurosci. Methods 53, 55–63. doi:10.1016/0165-0270(94)90144-9.

Charles, J. P., Cappellari, O., Spence, A. J., Hutchinson, J. R., and Wells, D. J. (2016). Musculoskeletal geometry, muscle architecture and functional specialisations of the mouse hindlimb. PLoS One 11, e0147669. doi:10.1371/journal.pone.0147669.

Christie, B. P., Freeberg, M., Memberg, W. D., Pinault, G. J. C., Hoyen, H. A., Tyler, D. J., and Triolo, R. J. (2017). “Long-term stability of stimulating spiral nerve cuff electrodes on human peripheral nerves”. J Neuroeng Rehabil 14, 70. doi:10.1186/s12984-017-0285-3.

Cloutier, S., LaFollette, M. R., Gaskill, B. N., Panksepp, J., and Newberry, R. C. (2018). Tickling, a technique for inducing positive affect when handling rats. J. Vis. Exp. doi:10.3791/57190.

Cotter, M., Cameron, N. E., Lean, D. R., and Robertson, S. (1989). Effects of long-term streptozotocin diabetes on the contractile and histochemical properties of rat muscles. Q. J. Exp. Physiol. 74, 65–74. doi:10.1113/expphysiol.1989.sp003240.

Fey, A., Schachner, M., and Irintchev, A. (2010). A novel motion analysis approach reveals late recovery in C57BL/6 mice and deficits in NCAM-deficient mice after sciatic nerve crush. J. Neurotrauma 27, 815–828. doi:10.1089/neu.2009.1217.

Fishman, M. A., Antony, A., Esposito, M., Deer, T., and Levy, R. (2019). The Evolution of Neuromodulation in the Treatment of Chronic Pain: Forward-Looking Perspectives. Pain Med. 20, S58–S68. doi:10.1093/pm/pnz074.

Forbes, J. M., and Cooper, M. E. (2013). Mechanisms of diabetic complications. Physiol. Rev. 93, 137–188. doi:10.1152/physrev.00045.2011.

Gamble, P., Stephen, M., MacEwan, M., and Ray, W. Z. (2016). Serial assessment of functional recovery following nerve injury using implantable thin-film wireless nerve stimulators. Muscle Nerve 54, 1114–1119. doi:10.1002/mus.25153.

Geremia, N. M., Gordon, T., Brushart, T. M., Al-Majed, A. A., and Verge, V. M. K. (2007). Electrical stimulation promotes sensory neuron regeneration and growth-associated gene expression. Exp. Neurol. 205, 347–359. doi:10.1016/j.expneurol.2007.01.040.

Gordon, T., Amirjani, N., Edwards, D. C., and Chan, K. M. (2010). Brief post-surgical electrical stimulation accelerates axon regeneration and muscle reinnervation without affecting the functional measures in carpal tunnel syndrome patients. Exp. Neurol. 223, 192–202. doi:10.1016/j.expneurol.2009.09.020.

Gordon, T., and English, A. W. (2016). Strategies to promote peripheral nerve regeneration: electrical stimulation and/or exercise. Eur. J. Neurosci. 43, 336–350. doi:10.1111/ejn.13005.

Günter, C., Delbeke, J., and Ortiz-Catalan, M. (2019). Safety of long-term electrical peripheral nerve stimulation: review of the state of the art. J Neuroeng Rehabil 16, 13. doi:10.1186/s12984-018-0474-8.

Isaacs, J., Mallu, S., Yan, W., and Little, B. (2014). Consequences of oversizing: nerve-to-nerve tube diameter mismatch. J. Bone Joint Surg. Am. 96, 1461–1467. doi:10.2106/JBJS.M.01420.

Jaiswal, M., Divers, J., Dabelea, D., Isom, S., Bell, R. A., Martin, C. L., Pettitt, D. J., Saydah, S., Pihoker, C., Standiford, D. A., et al. (2017). Prevalence of and risk factors for diabetic peripheral neuropathy in youth with type 1 and type 2 diabetes: SEARCH for diabetes in youth study. Diabetes Care 40, 1226–1232. doi:10.2337/dc17-0179.

Koo, J., MacEwan, M. R., Kang, S.-K., Won, S. M., Stephen, M., Gamble, P., Xie, Z., Yan, Y., Chen, Y.-Y., Shin, J., et al. (2018). Wireless bioresorbable electronic system enables sustained nonpharmacological neuroregenerative therapy. Nat. Med. 24, 1830–1836. doi:10.1038/s41591-018-0196-2.

Kruspe, M., Thieme, H., Guntinas-Lichius, O., and Irintchev, A. (2014). Motoneuron regeneration accuracy and recovery of gait after femoral nerve injuries in rats. Neuroscience 280, 73–87. doi:10.1016/j.neuroscience.2014.08.051.

Kumar, K., Taylor, R. S., Jacques, L., Eldabe, S., Meglio, M., Molet, J., Thomson, S., O’Callaghan, J., Eisenberg, E., Milbouw, G., et al. (2008). The effects of spinal cord stimulation in neuropathic pain are sustained: a 24-month follow-up of the prospective randomized controlled multicenter trial of the effectiveness of spinal cord stimulation. Neurosurgery 63, 762–70; discussion 770. doi:10.1227/01.NEU.0000325731.46702.D9.

Lin, Y.-C., Kao, C.-H., Chen, C.-C., Ke, C.-J., Yao, C.-H., and Chen, Y.-S. (2015). Time-course effect of electrical stimulation on nerve regeneration of diabetic rats. PLoS One 10, e0116711. doi:10.1371/journal.pone.0116711.

Lozano, A. M., Lipsman, N., Bergman, H., Brown, P., Chabardes, S., Chang, J. W., Matthews, K., McIntyre, C. C., Schlaepfer, T. E., Schulder, M., et al. (2019). Deep brain stimulation: current challenges and future directions. Nat. Rev. Neurol. 15, 148–160. doi:10.1038/s41582-018-0128-2.

MacEwan, M. R., Gamble, P., Stephen, M., and Ray, W. Z. (2018). Therapeutic electrical stimulation of injured peripheral nerve tissue using implantable thin-film wireless nerve stimulators. J. Neurosurg., 1–10. doi:10.3171/2017.8.JNS163020.

Martin, L. M., and McCabe, L. R. (2007). Type I diabetic bone phenotype is location but not gender dependent. Histochem. Cell Biol. 128, 125–133. doi:10.1007/s00418-007-0308-4.

McCrery, R., Lane, F., Benson, K., Taylor, C., Padron, O., Blok, B., De Wachter, S., Pezzella, A., Gruenenfelder, J., Pakzad, M., et al. (2020). Treatment of Urinary Urgency Incontinence Using a Rechargeable SNM System: 6-Month Results of the ARTISAN- SNM Study. J. Urol. 203, 185–192. doi:10.1097/JU.0000000000000458.

Melendez-Ramirez, L. Y., Richards, R. J., and Cefalu, W. T. (2010). Complications of type 1 diabetes. Endocrinol Metab Clin North Am 39, 625–640. doi:10.1016/j.ecl.2010.05.009.

Mobbs, R. J., Nair, S., and Blum, P. (2007). Peripheral nerve stimulation for the treatment of chronic pain. J. Clin. Neurosci. 14, 216–21; discussion 222. doi:10.1016/j.jocn.2005.11.007.

Morrow, T. J. (2004). Animal models of painful diabetic neuropathy: the STZ rat model. Curr Protoc Neurosci Chapter 9, Unit 9.18. doi:10.1002/0471142301.ns0918s29.

Mosconi, T., and Kruger, L. (1996). Fixed-diameter polyethylene cuffs applied to the rat sciatic nerve induce a painful neuropathy: ultrastructural morphometric analysis of axonal alterations. Pain 64, 37–57. doi:10.1016/0304-3959(95)00077-1.

Onode, E., Uemura, T., Takamatsu, K., Shintani, K., Yokoi, T., Okada, M., and Nakamura, H. (2019). Nerve capping with a nerve conduit for the treatment of painful neuroma in the rat sciatic nerve. J. Neurosurg., 1–9. doi:10.3171/2018.10.JNS182113.

Reardon, S. (2014). Electroceuticals spark interest. Nature 511, 18. doi:10.1038/511018a.

Rutschmann, M., Dahlmann, B., and Reinauer, H. (1984). Loss of fast-twitch isomyosins in skeletal muscles of the diabetic rat. Biochem. J. 221, 645–650. doi:10.1042/bj2210645.

Scheller, E. L., Cawthorn, W. P., Burr, A. A., Horowitz, M. C., and MacDougald, O. A. (2016). Marrow adipose tissue: trimming the fat. Trends Endocrinol. Metab. 27, 392–403. doi:10.1016/j.tem.2016.03.016.

Silva, M. J., Brodt, M. D., Lynch, M. A., McKenzie, J. A., Tanouye, K. M., Nyman, J. S., and Wang, X. (2009). Type 1 diabetes in young rats leads to progressive trabecular bone loss, cessation of cortical bone growth, and diminished whole bone strength and fatigue life. J. Bone Miner. Res. 24, 1618–1627. doi:10.1359/jbmr.090316.

Singh, B., Krishnan, A., Micu, I., Koshy, K., Singh, V., Martinez, J. A., Koshy, D., Xu, F., Chandrasekhar, A., Dalton, C., et al. (2015). Peripheral neuron plasticity is enhanced by brief electrical stimulation and overrides attenuated regrowth in experimental diabetes. Neurobiol. Dis. 83, 134–151. doi:10.1016/j.nbd.2015.08.009.

Tomlinson, R. E., Li, Z., Zhang, Q., Goh, B. C., Li, Z., Thorek, D. L. J., Rajbhandari, L., Brushart, T. M., Minichiello, L., Zhou, F., et al. (2016). NGF-TrkA Signaling by Sensory Nerves Coordinates the Vascularization and Ossification of Developing Endochondral Bone. Cell Rep. 16, 2723–2735. doi:10.1016/j.celrep.2016.08.002.

Wee, N. K. Y., Lorenz, M. R., Bekirov, Y., Jacquin, M. F., and Scheller, E. L. (2019). Shared autonomic pathways connect bone marrow and peripheral adipose tissues across the central neuraxis. Front. Endocrinol. (Lausanne) 10, 668. doi:10.3389/fendo.2019.00668.

Wong, J. N., Olson, J. L., Morhart, M. J., and Chan, K. M. (2015). Electrical stimulation enhances sensory recovery: a randomized controlled trial. Ann. Neurol. 77, 996–1006. doi:10.1002/ana.24397.

Xu, J., Wang, J., Chen, X., Li, Y., Mi, J., and Qin, L. (2020). The Effects of Calcitonin Gene-Related Peptide on Bone Homeostasis and Regeneration. Curr. Osteoporos. Rep. doi:10.1007/s11914-020-00624-0.

Yao, G., Kang, L., Li, J., Long, Y., Wei, H., Ferreira, C. A., Jeffery, J. J., Lin, Y., Cai, W., and Wang, X. (2018). Effective weight control via an implanted self-powered vagus nerve stimulation device. Nat. Commun. 9, 5349. doi:10.1038/s41467-018-07764-z.

