## Supplemental Material for "Neuroskeletal effects of chronic bioelectric nerve stimulation in health and diabetes"

### *Supplementary Material*

**Supplementary Video 1.** Video recording of rat undergoing bioelectric nerve stimulation treatment. External transmitter coil was placed over the implanted coil. Each “beep” corresponds to an increase in stimulus amplitude. End of movie illustrates “maximally-activated effect” mentioned in Methods with pointed toes and clenched paw. This position was sustained for the one-hour treatment regimen.
